# ATR and mTOR regulate F-actin to alter nuclear architecture and repair replication stress

**DOI:** 10.1101/451708

**Authors:** Noa Lamm, V. Pragathi Masamsetti, Mark N. Read, Maté Biro, Anthony J. Cesare

## Abstract

“Replication stress” describes phenomena that alter DNA replication rates ^1–3^. Multiple architectural challenges within the confined nuclear volume must be navigated during replication to prevent or repair replication stress. Cellular mechanisms potentiating changes in nuclear architecture that facilitate DNA replication remain unclear. Here we show that the ATR, IPMK and mTOR kinases regulate actin polymerisation in human cells to alter nuclear architecture and promote replication fork repair. We demonstrate that replication stress activates mTOR, in an ATR and IPMK-dependent manner, to induce polymerisation of nuclear filamentous actin (F-actin). mTOR and ATR then counteract replication stress-induced nuclear envelope deformation and increase nuclear volume through their regulation of actin dynamics. Additionally, we reveal that FANCD2 labelled replication forks colocalise with actin filaments in late S-phase. mTOR and ATR then regulate the mobility, speed and directionality of stalled replication foci within the three-dimensional nuclear architecture. Importantly, we find nuclear F-actin also acts as a substrate for the directed migration of stalled replication foci to the nuclear periphery. Suppressing mTOR and ATR-dependent actin forces prevents replication fork restart and promotes chromosome segregation errors in primary and cancer cell lines. Together, these data reveal that ATR and mTOR regulate actin dynamics in the replication stress response to alter nuclear architecture and maintain genome stability.

## MAIN TEXT

Actin is a cytoskeletal protein with key roles in cargo transport and modulation of cellular shape and motility ^4^. These functions are mainly regulated via spatiotemporally-controlled polymerisation of monomeric actin into filamentous actin (F-actin). Although actin is largely perceived as a cytoplasmic protein, transient F-actin is observable within eukaryotic nuclei ^5^, where it is involved in a variety of cellular processes such as the serum response ^6^, cell spreading ^7^, mitotic exit ^8^ and DNA repair ^9–11^.

During DNA replication, cells must navigate diverse architectural challenges. This includes mediating passage of DNA polymerases across the entirety of the genome; coping with nuclear envelope deformation generated by torsional stress in chromatin attached to the nuclear periphery ^12^; and coordinating the stability and repair of stalled replication forks throughout the nuclear volume ^13^. It was suggested that actin dynamics in G1 promote replication initiation through transcriptional regulation ^14^. However, it is unknown if actin-dependent forces contribute to facilitating effective DNA replication during S-phase.

To determine if actin polymerisation impacts DNA replication in human cells, we treated IMR90 primary fibroblasts, IMR90 fibroblasts expressing HPV 16 E6 and E7 (IMR90 E6E7), and U-2OS osteosarcoma cells with the actin polymerisation inhibitor Latrunculin B (LatB) ^15^, in the presence or absence of the Family B DNA polymerase inhibitor Aphidicolin (APH). Consistent with replication stress, DNA fibre assays revealed that LatB induced a significant reduction in replication fork speed and fork distance, which was additive with APH co-treatment (Fig. 1a-c, Extended Data Fig.1). In agreement, live cell imaging of LatB treated *f*luorescent, *u*biquitination-based *c*ell *c*ycle *i*ndicator (FUCCI) ^16^ modified IMR90 E6E7 cells revealed an extended S-phase duration, indicative of replication stress (Fig. 1d).

**Fig. 1:**
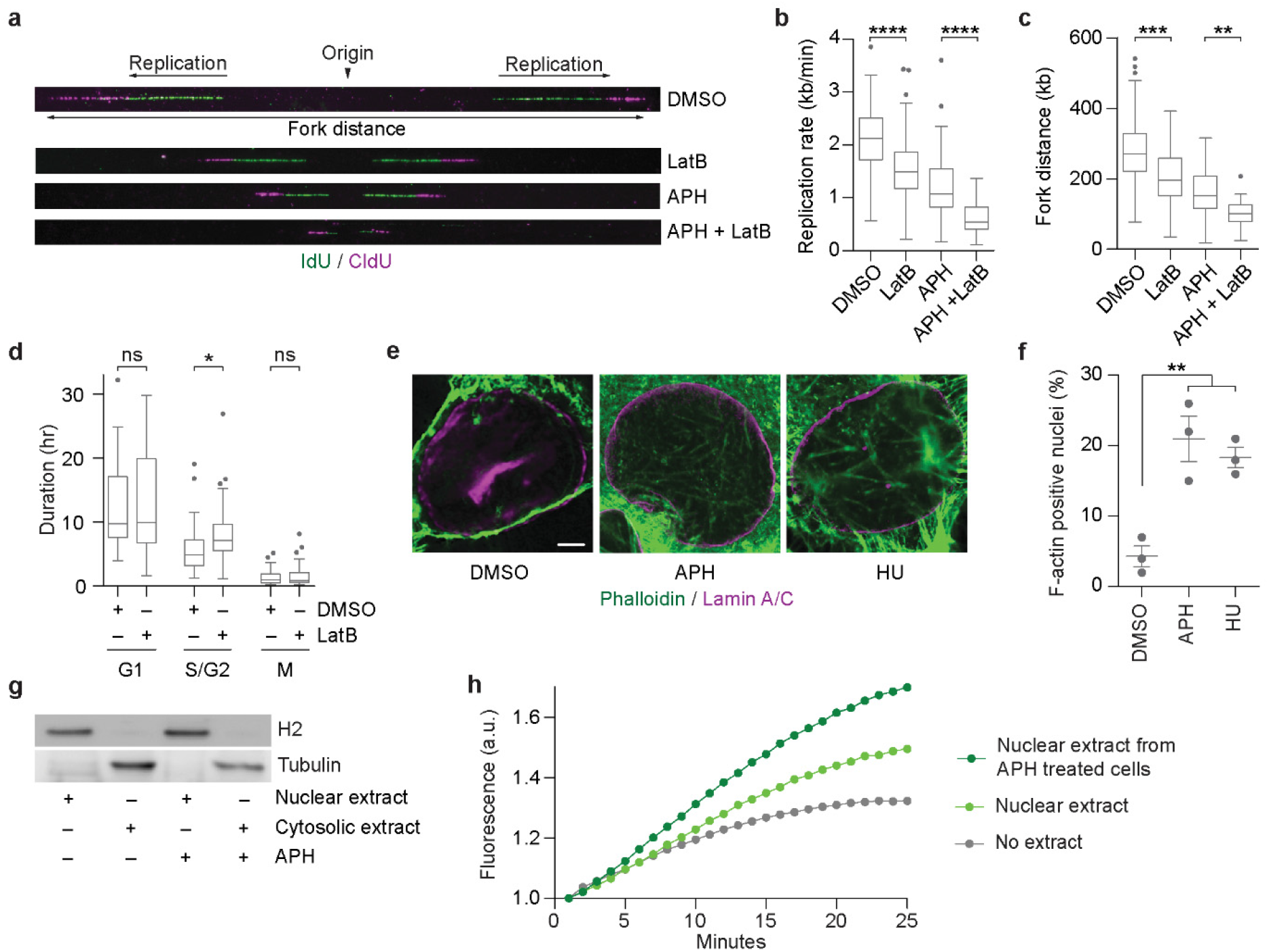
Replication stress induces nuclear actin polymerisation. **a,**Representative images of chromatin fibre assays from IMR90 cells treated with 0.01 μM Aphidicolin (APH) and/or 0.2 μM Latrunculin B (LatB) for two hours (one hour before and one hour during pulse-labelling with thymidine analogous). Replication rates were calculated from IdU track length. Replication fork distance is the length from the ends of the two CldU tracks. **b-c,** Replication rate **b** and distance **c** from the experiment depicted in **a** (n > 130 forks per condition over three biological replicates compiled into a Tukey box plot, student’s t-test). **d,** Cell cycle phase duration from IMR90 E6E7-FUCCI cells treated with 0.2 μM LatB (n > 42 cells over three biological replicates compiled into a Tukey box plot, two-tailed student’s t-test). **e,** Representative super resolution image of a single Z-plane from U-2OS cells stained with phalloidin and α-Lamin A/C following treatment with DMSO, 0.4 μM APH, or 500 μM hydroxyurea (HU) for 24 hours. Scale bar represents 5 μm. **f,** Frequency of nuclear actin fibre positive nuclei from experiments shown in **e** (mean ± s.e.m, each data point represents an individual biological replicate, n > 99 nuclei per condition over three biological replicates, Fisher’s exact test). **g,** Western blot of nuclear and cytosolic extracts prepared from IMR90 cells ± 0.4 μM APH treatment for eight hours. **h,** Representative normalised time course of pyrene-labelled actin assembly in the presence or absence of IMR90 nuclear extracts from **g.** One of three independent biological replicates are shown. For all panels: ns = not significant, *\*p < 0.05, **p < 0.005, ***p < 0.001, ****p < 0.0001*.

To determine whether inducing DNA replication stress impacted actin dynamics, we treated U-2OS cells with APH or the ribonucleotide reductase inhibitor Hydroxyurea (HU). Actin fibres were labeled with phalloidin and imaged with super-resolution microscopy. We found that replication stress promoted F-actin formation in human nuclei (Fig. 1e, f). To specifically visualise the low-abundance pool of nuclear actin, we expressed an actin chromobody fused on its N-terminus with a nuclear localisation signal (NLS) and GFP (NLS-GFP-actin-CB) ^7^. In agreement with phalloidin staining, the percentage of F-actin positive nuclei in NLS-GFP-actin-CB transfected IMR90, IMR90 E6E7, and U-2OS cells all significantly increased with APH or HU treatment (Extended Data Fig. 2a-d). In vitro pyrene actin assays ^6^ also revealed that actin polymerisation rates increased in nuclear extracts from IMR90, IMR90 E6E7 and U-2OS cells when cultures were pre-treated with APH (Fig. 1g, h, Extended Data Fig. 2e-h). Together, these data indicate that replication stress induces nuclear F-actin formation and that inhibiting actin polymerisation impedes DNA replication in a variety of human primary and cancer cells.

**Fig. 2:**
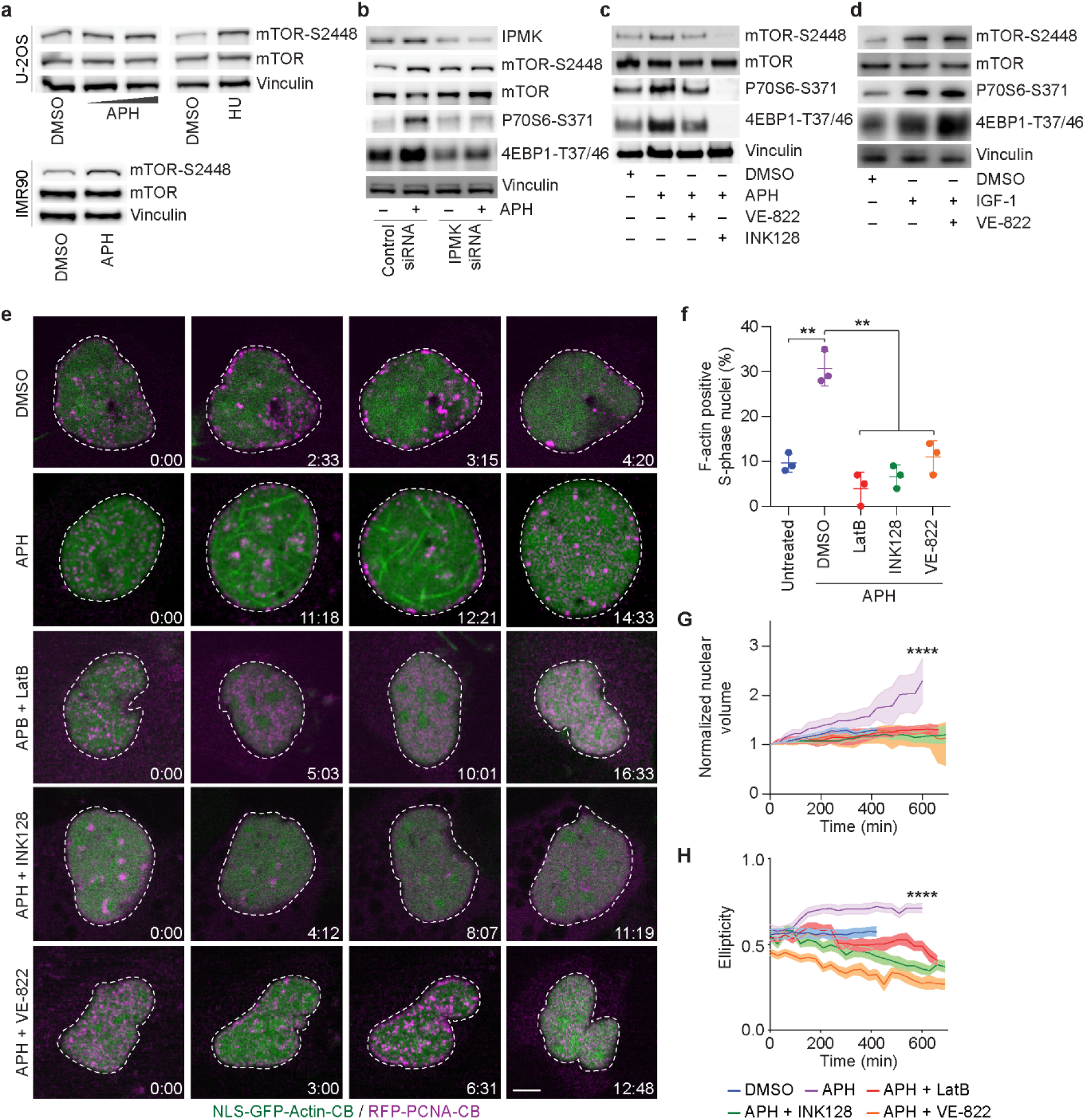
ATR and mTOR regulate F-actin dependent changes to nuclear architecture in response to replication stress. **a,** Western blots of whole cell extracts from U-2OS and IMR90 cells treated with 0.4 μM APH or 500 μM HU for eight hours. **b,** Western blots of whole cell extracts from siRNA transfected HT1080 6TG cells treated with or without 0.4 μM APH for eight hours. Cells were siRNA transfected 48 hours prior to extraction. **c,** Western blots of whole cell extracts from HT1080 6TG cells treated with 0.4 μM APH for eight hours ± 0.2 μM INK128 or 1 μM VE-822. **d,** Western blots of whole cell extracts from HT1080 6TG cells treated with 100 ng/ml IGF-1 ± 1 μM VE-822 for one hour. **e,** Representative images of spinning-disk confocal live cell microscopy of U-2OS cells transfected with NLS-GFP-actin-CB and RFP-PCNA-CB, treated with 0.4 μM APH ± 0.2 μM LatB, 0.2 μM INK128, or 1 μM VE-822. Time is shown as (hr:min) relative to the first image of the series. Scale bar represents 5 μm. **f,** Frequency of nuclear F-actin positive S-phase nuclei from the images depicted in e (mean ± s.e.m, each data point represents an individual biological replicate, n > 200 nuclei per condition over three biological replicates, Fisher’s exact test). **g-h,** Time courses of normalised nuclear volume g and ellipticity h for the experiment images depicted in e (mean ± s.e.m., n > 31 nuclei per condition over five biological replicates, one-way ANOVA). For all panels: *\*\*p < 0.01, ****p < 0.0001*.

We predicted that the ataxia telangiectasia mutated and Rad3-related (ATR), inositol polyphosphate multikinase (IPMK), and mammalian target of rapamycin (mTOR) kinases functioned as regulators of replication stress-induced actin dynamics. This is because ATR is the master regulator of the replication stress response ^17^ and is activated by double strand breaks in a pathway involving IPMK and nuclear actin polymerisation ^18^. Additionally, mTOR, which functions downstream of IPMK, is a regulator of actin polymerisation, and has an essential but unidentified role in the replication stress response ^19^. Accordingly, we found in a variety of human primary and cancer cells that APH or HU treatment increased mTOR-S2448 phosphorylation, indicative of elevated mTOR activity (Fig. 2a-c, Extended Data Fig. 3a) ^20^. Replication stress-induced mTOR activity was ATR and IPMK-dependent, as IPMK depletion or ATR inhibition with VE-822 ^21^ both suppressed elevated phosphorylation of mTOR and its downstream targets P70S6 and 4EBP in response to APH (Fig. 2b, c, Extended Data Fig. 3). Inhibiting mTOR with INK128 ^22^ completely suppressed the mTOR signaling pathway (Fig. 2c, Extended Data Fig. 3a). Notably, VE-822 did not suppress mTOR activation by IGF-1 ^23^ (Fig. 2d), indicating that ATR regulates mTOR specifically in response to replication stress. Additionally, INK128 did not suppress CHK1 phosphorylation by ATR, indicating that IPMK and ATR are upstream of mTOR in the replication stress response (Extended Data Fig. 3a).

To visualise nuclear actin dynamics specifically in cells undergoing DNA replication, we cotransfected IMR90 and U-2OS cells with the NLS-GFP-actin-CB and an RFP tagged chromobody against PCNA (RFP-PCNA-CB). RFP-PCNA-CB foci identify the sites of active DNA replication and serve as a marker of S-phase. We treated asynchronous cultures with APH ± DMSO vehicle, LatB, VE-822 or INK128, and imaged fixed (Extended Data Fig. 4a) and live cells (Fig. 2e, Videos 1-4). APH treatment induced a significant increase in nuclear F-actin networks specifically during S-phase, dependent on actin polymerisation, and ATR and mTOR activity (Fig. 2f, Extended Data Fig. 4b, c). Depleting ATR or mTOR likewise suppressed S-phase nuclear F-actin in APH treated cells (Extended Data Fig. 4d, e). Inhibiting mTOR or ATR also induced a striking phenotype in some APH treated cells, where nuclear F-actin assembled for a brief time, before losing its structural integrity (Extended Data Fig. 4f, Video 5).

Fixed and live-cell imaging revealed substantial F-actin dependent changes to nuclear architecture accompanying replication stress. Notably, actin polymerisation expanded the nuclear volume when nuclear F-actin was present in APH treated S-phase cells (Fig. 2e, g, Extended Data Fig. 5, Videos 2, 3). Consistent with ATR and mTOR regulation, this increased nuclear volume was suppressed with VE-822, INK128, and mTOR or ATR siRNAs (Fig. 2e, g, Extended Data Fig. 5b, c, Video 4). Actin polymerisation also significantly increased nuclear ellipticity following replication stress in an mTOR and ATR dependent manner, as demonstrated by phenotype suppression with LatB, INK128 or VE-822 (Fig. 2e, h). Correspondingly, inhibiting actin polymerisation in APH treated cells resulted in nuclear invagination and deformation (Fig. 2e, Extended Data Fig. 4a). mTOR and ATR thus regulate F-actin polymerisation to shape nuclear architecture and counteract replication stress-induced nuclear malformation ^12^.

Analysing the position of RFP-PCNA-CB foci within the three-dimensional nuclear volume of fixed IMR90 cells revealed that replication foci localise with greater frequency to the nuclear periphery in APH treated cultures (Extended Data Fig. 6a, b). Moreover, RFP-PCNA-CB localisation to the nuclear exterior was dependent on actin polymerisation, and ATR and mTOR activity (Extended Data Fig. 6a, b). Fixed images revealed that the “difficult to replicate” telomeric DNA ^24^ also localised towards the nuclear periphery in an actin polymerisation dependent manner in APH treated cells (Extended Data Fig. 6c, d). Together these data suggested that actin polymerisation mobilised stressed replication foci in human cells.

To test this directly, we sought to track RFP-PCNA-CB foci in live cell imaging experiments. In these experiments we focused on late replication foci for two reasons. First, late replication foci are present after heterochromatic regions at the nuclear periphery are synthesized ^25^, which removes the potential bias of analysing foci that originate at the nuclear periphery in mid S-phase (Extended Data Fig. 7a). Second, late replication foci in APH treated cells display extensive colocalisation with FANCD2, indicative of stalled replication forks ^26^ (Fig. 3a). Notably, we observed that colocalised FANCD2 and RFP-PCNA-CB foci also associated with actin fibres in super resolution images of late S-phase cells (Fig. 3a).

**Fig. 3:**
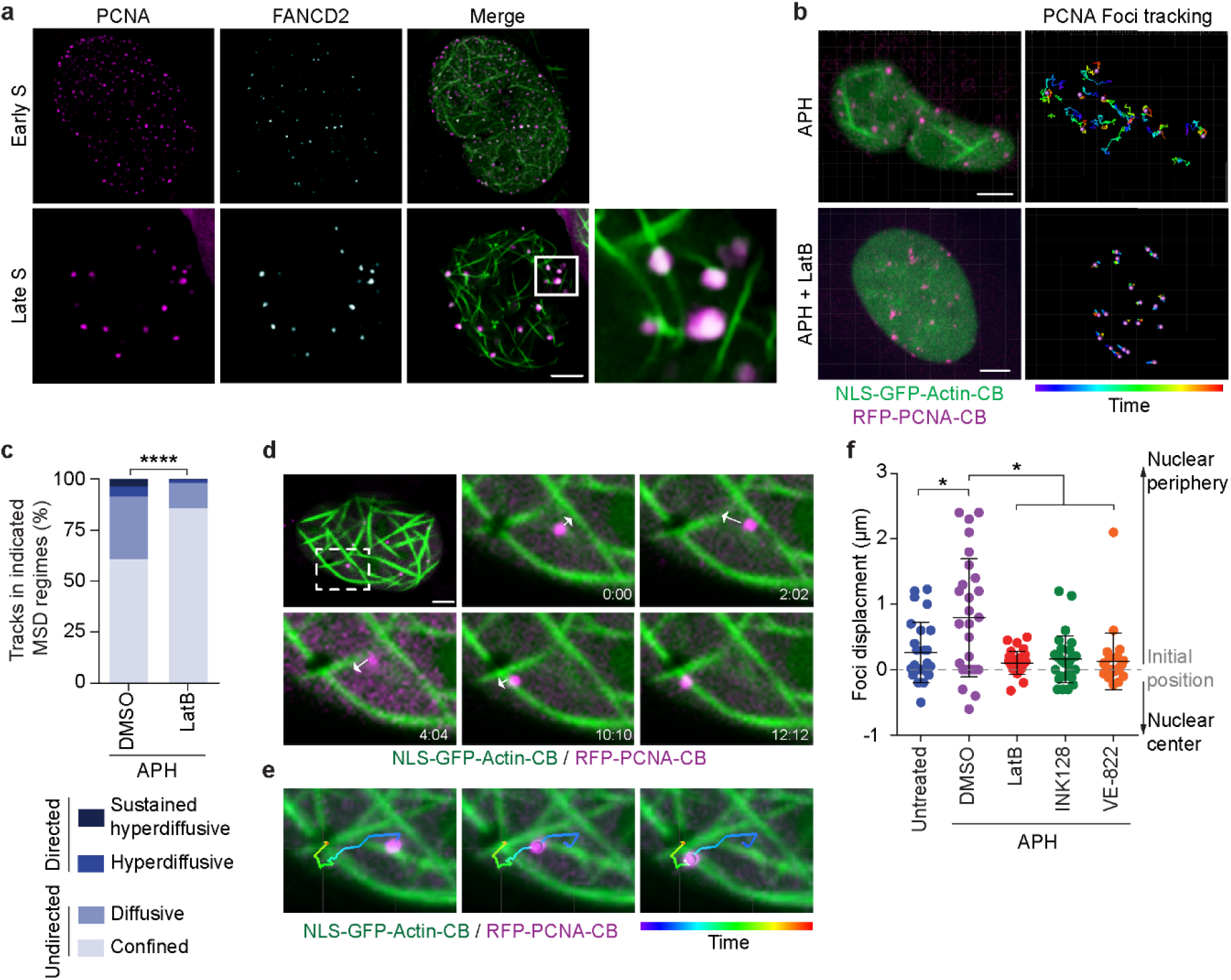
Nuclear F-actin mediates directed movement of stalled late replication foci towards the nuclear periphery. **a,** Representative super resolution images of a single Z-plane from NLS-GFP-actin-CB and RFP-PCNA-CB transfected U-2OS cells treated with 0.4 μM APH for 24 hours. Cells were fixed and stained with FANCD2 immunofluorescence. **b,** Left: Representative maximum projection images captured from spinning-disk confocal live cell microscopy of NLS-GFP-actin-CB and RFP-PCNA-CB transfected U-2OS cells treated with 0.4 μM APH ± 0.2 μM LatB. Right: RFP-PCNA-CB foci tracks from the video on the left. **c,** Percentage of late S phase RFP-PCNA-CB foci tracks spent in the indicated MSD regimes in U-2OS cells treated with 0.4 μM APH ± 0.2 μM LatB (n > 45 tracks from at least four nuclei for each condition, Chi-squared test). **d,** Representative single Z plane from spinning-disk confocal live cell microscopy of NLS-GFP-actin-CB and RFP-PCNA-CB transfected U-2OS cells treated with 0.4 μM APH. Time is shown as (min:sec) relative to the first image of the series. Arrows indicate observed foci movement. **e,** Automated foci tracking in the images depicted in **d. f,** Direction of late replication foci displacement in NLS-GFP-actin-CB and RFP-PCNA-CB transfected U-2OS cells treated with 0.4 μM APH ± 0.2 μM LatB, 0.2 μM INK128, or 1 μM VE-822 (n > 26 late S foci from at least seven nuclei, Kruskal-wallis test). For all panels: Scale bar represents 5 μm and *\*p < 0.05, ****p < 0.0001*.

To track foci migration, we developed image analysis tools that register nucleus location in progressive live-cell imaging frames to correct for natural cell motility (Video 6). Individual late S-phase RFP-PCNA-CB foci were then tracked from APH treated U-2OS cells in the presence or absence of LatB, INK128 or VE-822, and the foci speed, mean-squared displacement (MSD), and total displacement calculated (Fig. 3b-f, Extended Data Fig. 7b-d, Videos 7-9). Replication stress increased the average speed of late replication foci movement within the nuclear volume, dependent upon actin polymerisation, and ATR and mTOR activity (Extended Data Fig. 7b). Inhibiting actin polymerisation also constrained foci trajectories in APH treated cells (Fig. 3b, Videos 7, 8).

To study the potentially heterogeneous phases of motion of late replication foci, we performed MSD analysis and classified foci movements into confined, diffusive, hyperdiffusive and sustained hyperdiffusive regimes. Detailed examination of individual RFP-PCNA-CB foci tracks in APH treated cells revealed bursts of sustained hyperdiffusive movements (Extended Data Fig. 7c). Strikingly, replication foci no longer displayed these sustained directional movements after inhibiting actin polymerisation in APH treated cells, and instead predominantly displayed confined motion (Fig. 3c, Extended Data Fig. 7c, d). Visualising individual RFP-PCNA-CB foci in APH treated cells revealed directed movement of RFP-PCNA-CB foci along actin fibres (Fig. 3d, e, Video 9). Quantifying the directionality of RFP-PCNA-CB foci mobility revealed that late replication foci in APH treated cells moved toward the nuclear periphery and did so in a manner dependent upon actin polymerisation, and ATR and mTOR activity (Fig. 3f). These data reveal that nuclear actin filaments enable increased diffusive movement of damaged replication foci as well as facilitate the directed movement of those forks to the nuclear periphery.

DSBs were proposed to accumulate at the nuclear periphery to finalise homologous recombination ^27^. Additionally, nuclear membrane associated lamin proteins were suggested to function in repair of stalled replication forks ^28^. We therefore hypothesised that F-actin dependent localisation of stalled replication forks to the nuclear periphery is required for fork repair. To test this, U-2OS and IMR90 cells were treated with APH for three hours, followed by APH washout and recovery in media containing vehicle, LatB, VE-822 or INK128 (Fig. 4a). Chromatin fibre assays identified that inhibiting F-actin network formation with LatB, as well as ATR or mTOR inhibition, suppressed replication fork restart (Fig. 4b, c, Extended Data Fig. 8a, b). Additionally, ATR signaling remained active in IMR90 E6E7 cells after APH washout if actin polymerisation was inhibited during the recovery period with LatB (Fig. 4d, e). A failure to repair stalled replication forks is expected to confer genome instability ^29^. In agreement, treating IMR90 or U-2OS cells with escalating dosages of LatB induced concomitant increases of micronuclei and anaphase abnormalities consistent with unrepaired replication stress (Fig. 4f, g, Extended Data Fig. 8c, d). Genome instability in LatB treated cultures was additive when the cells were co-treated with APH (Fig. 4f, g, Extended Data Fig. 8c, d).

**Fig. 4:**
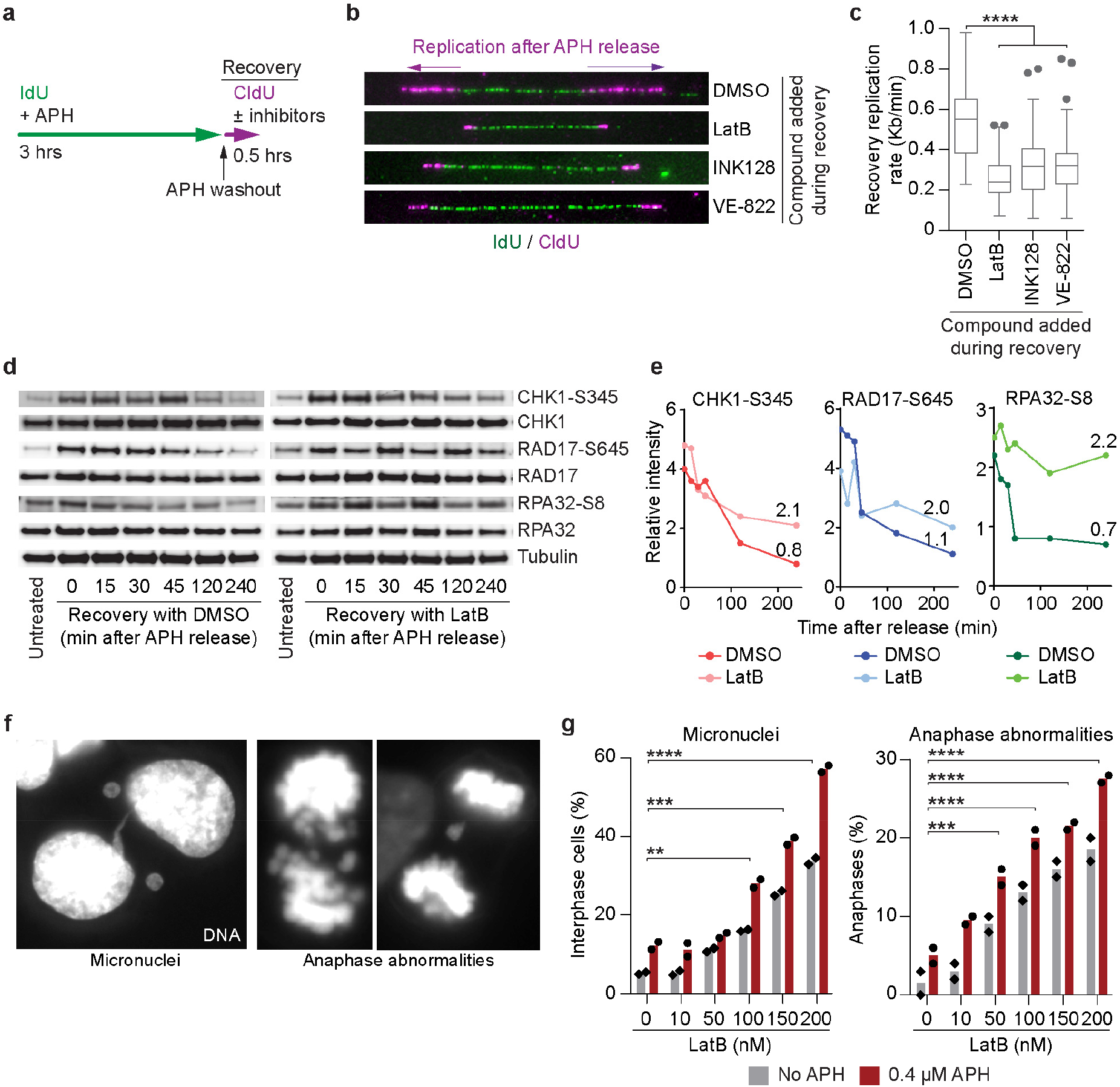
Nuclear F-actin mediates replication fork repair to maintain genome stability. **a,** Illustration of the experimental scheme in **b. b,** Representative images of chromatin fibre assays to measure the efficiency of replication fork restart in U-2OS cells after APH washout in the presence of DMSO, 0.2 μM LatB, 0.2 μM INK128, or 1 μM VE-822. Replication rate is measured from the CldU track. c, Quantitation of the experiment shown in **b** (n > 132 replication forks from two independent experiments compiled into a Tukey box plot, two-tailed student’s t-test). **d,** Western blots of whole cell extracts from IMR90 E6E7 cells harvested at the indicted time points following release from 24 hours of 0.4 μM APH treatment ± 0.2 μM LatB. Representative blots from one of three biological replicates are shown. e, Signal intensity at the indicated time points from the blots shown in **d,** relative to the untreated extract. One of three independent biological replicates are shown. f, Representative images of micronuclei (left panel) and anaphase abnormalities (middle and right panels) in U-2OS cells following treatment with escalating dosages of LatB. g, Frequency of micronuclei (left) and anaphase abnormalities (right) from the images depicted in **f** (n ≥ 152 cells for each concentration over two biological replicates, individual replicate means are shown by the data points, bar represents the overall mean. Fisher’s exact test). For all panels: ns = not significant, *\*\*p < 0.001, ***p < 0.0005, ****p < 0.0001*.

Together, our data identify a pathway regulated by ATR, IPMK and mTOR, which regulates actin polymerisation to shape nuclear architecture and to promote replication fork repair. ATR and IPMK are co-regulated by genomic damage ^18^ and our data indicate that these kinases function upstream of mTOR during replication stress. Our data also reveal the uncharacterised role for mTOR in the replication stress response^19^ to include regulation of actin dynamics. We suggest ATR and mTOR alter nuclear architecture through forces potentiated by nuclear F-actin. This is supported by the specific induction of nuclear F-actin in S-phase cells under replication stress; the temporal correlation between nuclear volume expansion and nuclear F-actin; and the directed movement of stalled replication foci along nuclear actin fibres. Our interpretation is that enhanced motility of stalled replication foci facilitates replication fork restart. Homologous recombination is promoted through chromatin mobility ^30^, and nuclear pores and the nuclear lamina are implicated in homologous recombination and replication fork repair ^28,31^. Inhibiting actin polymerisation simultaneously confined replication foci movement, inhibited fork repair, and promoted genome instability: observations consistent with F-actin mobilising stalled forks for repair. During the course of our study, two independent reports identified that F-actin promotes double strand break repair^10,11^. Data presented here substantially expand our understanding of nuclear F-actin in genome stability by identifying its critical role in the replication stress response, elucidating the underlying signaling pathways controlling actin-dependent repair, and demonstrating the role of F-actin in replication foci mobility and modulation of nuclear architecture. Together these concerted discoveries reveal the extensive role for F-actin in genome maintenance.

## METHODS

### Cell culture and treatments

IMR90, IMR90 E6E7 ^32^, and U-2OS cells were provided by Jan Karlseder (Salk Institute, La Jolla California), and HT1080 6TG cells by Eric Stanbridge (University of California, Irvine). Cell line identity was verified by Cell Bank Australia using short tandem repeat profiling and all cell lines were identified as mycoplasma negative (MycoAlert, LT07-118, Lonza). All cells were grown at 37° C, 10% CO2, and 3% O2 in DMEM (Life Technologies) supplemented with 1% non-essential amino acids (Life Technologies), 1% Glutamax (Life Technologies) and 1% penicillin-streptomycin (Life Technologies). IMR90, IMR90 E6E7 and IMR90 E6E7-FUCCI were supplemented with 10% fetal bovine serum (Life Technologies), and U-2OS and HT1080 6TG cultures with 10% bovine growth serum (HyClone). The following compounds were used in cell treatments: dimethyl sulfoxide (DMSO, Sigma-Aldrich), Aphidicolin (APH, Sigma-Aldrich), Hydroxyurea (HU, Sigma-Aldrich), Latrunculin B (LatB, Cayman Chemical: 10010631), INK128 (Cayman Chemical: 11811), VE-822 (Selleckchem: S7102) and Insulinlike growth factor-I (IGF1, Sigma-Aldrich: SRP4121,).

### Chromatin fibre assays

Replication dynamics were measured using chromatin fibre analysis as described elsewhere ^33^. Briefly, unsynchronised cells were sequentially pulse-labelled with 100 μM of thymidine analogues IdU (Sigma) and then CldU (Sigma) for 30 min each. For experiments using chemical inhibitors, the compounds were added one hour prior to IdU addition and kept in the culture media through IdU and CldU labelling. After harvesting, the genomic DNA was stretched onto glass slides at a constant factor of 2 kb/μm using a molecular combing system (Genomic Vision). Nascent DNA replication was visualised by immunofluorescence against IdU and CldU and images captured using a ZEISS AxioImager Z.2, with a 63x 1.4 NA oil objective, appropriate filter cubes, an Axiocam 506 monochromatic camera (ZEISS), and Zen software (ZEISS). In our analysis, only replication forks with an origin of replication, and both IdU and CldU staining were scored. Replication rates were calculated solely on the IdU tracks, which result from the first pulse of nucleoside analogue. Fork distance is calculated as the length between two sister forks measured from the ends of the CldU tracks. Fork distance is approximately half the length of the entire replicon ^34^. Replicon length scales with increasing inter-origins distances and therefore serves as readout of the distance between activated origins ^34^. The length of replication signals and the distances between sister forks were converted to kilobasepairs according to a constant and sequence-independent stretching factor (1 μm = 2 Kb) using Zen software. As this assay is highly sensitive to replication stress-inducing agents, for these experiments we used 0.01 μM APH.

Replication fork restart was measured using chromatin fibre analysis as described above except that cells were pulse-labelled for three hours with IdU in the presence of 0.1 μM APH to arrest replication forks. APH and IdU were washed out of the culture and the cells treated for 30 min in the presence or absence of LatB, INK128 or VE-822. Restart efficiency was determined based on the length of the CldU tracks using the constant 1 μm = 2 Kb. The “n” for all chromatin fibre analysis describes the number of individual forks analysed. For distance analysis, this reflects half of the data points, one for two forks moving away from the origin.

### FUCCI live cell imaging and analysis

mVenus-hGeminin (1/110)/pCSII-EF and mCherry-hCdt1(30/120)/pCSII-EF were a kind gift from Atsushi Miyawaki ^16^ and IMR90 E6E7 FUCCI cells were created as described elsewhere ^35^. Cells were seeded 24 hours prior to imaging in 6-well glass bottom plates (World Precision Instruments). LatB was added two hours prior to imaging. Combinatorial fluorescence and Differential Interference Contrast microscopy were performed on a ZEISS Cell Observer inverted wide field microscope, with 20x 0.8 NA air objective, at 37° C, 10% CO2 and 3% oxygen. Images were captured every six minutes, for a duration of sixty hours, using an Axiocam 506 monochromatic camera and Zen software. Videos were scored by eye. Cell cycle was determined colourimetrically with G1 identified by mCherry-hCdt1(30/120) stability and S/G2 by mVenus-hGeminin (1/110) stability ^16^. Mitotic duration was determined by cell morphology.

### Phalloidin staining

Phalloidin staining was performed as described elsewhere ^6^. Briefly, cells were seeded on sterile glass cover slips 48 hours prior to fixation, and APH or HU were added to the culture media 24 hours prior to fixation. Initial fixation was done for 1 min in a mixture of 0.5% Triton X-100 and 0.25% glutaraldehyde (Sigma-Aldrich) in cytoskeleton buffer (10 mM MES, 150 mM NaCl, 5 mM EGTA, 5 mM glucose and 5 mM MgCl2 at pH 6.1). Post-fixation the coverslips were incubated for 15 min in cytoskeleton buffer containing 2% glutaraldeyde. Before phalloidin labeling, autofluorescence was quenched by treatment with 1 mg/ml freshly prepared sodium borohydride (Sigma-Aldrich). Cover slips were incubated overnight at 4° C with Phalloidin-Atto 488 (Sigma-Aldrich: 49409) according to manufacturer protocol. The following day, slides were washed four times, 5 min each in PBS and incubated for 3 hours at 37° C with anti-Lamin A/C antibody, washed four times in 1x PBS for 5 min each and then incubated with goat polyclonal anti-rabbit Alexa Fluor 568 secondary antibody for an hour in 37°. First and secondary antibodies were diluted in PBS supplemented with 5% FCS. Before mounting, slides were washed again as described above and dehydrated in a graded series (70, 90, 100%) of ethanol solutions for two min each. Slides were then mounted with Prolong Gold (Life Technologies) and imaged with super resolution microscopy as described below. To avoid complication of phalloidin signals arising from above or below the nucleus, imaging was limited to cells with a clearly resolvable nuclear interior.

### Pyrenyl-actin assembly assays

Nuclear extracts were prepared as described elsewhere ^6^. Briefly, when appropriate cells were treated with APH for eight hours. APH treated and untreated cells were then washed once with cold phosphate-buffered saline (PBS), scraped carefully, and centrifuged at 800g for 10 min. The cell pellet was kept at −80°C for 45 min, before resuspension in buffer P1 [10 mM Hepes, 0.1 mM EGTA, 1 mM DTT, complete protease inhibitors (Roche)]. After addition of Triton X100 to a final concentration of 0.5%, samples were vortexed for 10 s, followed by sedimentation of nuclei at 10,000g for 10 min. The nuclear pellet was washed with buffer P1 before lysis in buffer P2 (20 mM Hepes, 25% Glycerol, 400 mM NaCl, 1 mM EGTA, 1 mM DTT, complete protease inhibitors) for 90 min on a rotary shaker at 4°C. Remaining insoluble material was sedimented at 16,000g for 30 min. The purity of obtained extracts was controlled by immunoblotting for α-tubulin and H2 histone.

Pyrenyl-actin assays were adapted from previously described protocols ^36^. Nuclear extracts prepared as described above were dialysed with a mini Dialysis Unit (Thermofisher scientific) against XB buffer (10 mM Hepes, pH 7.7, 100 mM KCl, 2 mM MgCl2, 0.1 mM CaCl2, 5 mM EGTA, 1 mM dithiothreitol) for at least 3 hours. Nuclear extracts were used with pyrene-labeled actin from the Actin Polymerisation Biochem Kit (Cytoskeleton, Inc.: BK003) according to the manufacturer protocol. Pyrene fluorescence was measured at 407 nm with excitation at 365 nm, and the kinetics of actin filament assembly monitored using an EnSpire Multimode Plate Reader (PerkinElmer).

### Chromobody transfection and imaging

Plasmids expressing the NLS-GFP-actin chromobody (Chromotek: acg-n) or RFP-PCNA chromobody (Chromotek: ccr) were transfected using Lipofectamine LTX (Thermofisher Scientific), according to the manufacturer protocol. For fixed cell imaging, cells were seeded on sterile cover slips and transfected with chromobody expressing plasmids 24 hours later. Chemical inhibitors were added 48 hours after transfection for 24 hours. Cover slips were fixed in 3.7% formaldehyde/1x PBS for 10 minutes and permeabilised with 0.5% Triton/1xPBS for 10 minutes in room temperature. Slides were mounted with Prolong Gold (Life Technologies) and imaged with super resolution imaging as described below.

For live cell imaging, cells were seeded on three cm glass bottom dish (World Precision Instruments) and transfected with the chromobody plasmids 24 hours later. 48 hours post-transfection, cells were treated with the indicated compounds for 24 hours. The culture media was then replaced by colourless DMEM (FluoroBrite DMEM, Life Technologies) supplemented as described above and where applicable replenished with fresh APH, LatB, INK128 or VE-822. Imaging was performed on a Zeiss Cell Observer SD spinning disk confocal microscope. Cells were imaged using Differential Interference Contrast (DIC) microscopy combined with fluorescent imaging (7% excitation power of 561 nm laser, 1 x 1 binning, EM gain of 908 and 6.5% excitation power of 488 laser, 1 x 1 binning, EM gain of 527) using appropriate filter sets, a 63x 1.3 NA oil objective at 37° C, 10% CO2 and 3% oxygen. A total of 10 z-stacks (135 nm) were captured in an image scaled to 47504 × 37602 pixels at 10.05 μm × 7.96 μm. Images were captured with Zen software using an Evolve Delta (Photometrics) camera, every 90 – 200 seconds for up to 72 hours.

### Telomere detection in S-phase cells

Cells were grown on sterile glass cover slips and treated with the indicated compounds for 24 hours and then pulse labelled with 100 μM EdU (Invitrogen: C10339) for one hour. Cells were fixed in 3.7% formaldehyde/1x PBS for 15 min and permeabilised with 0.5% Triton/1x PBS for 20 min. EdU was detected using the Click-iT EdU imaging kit (Invitrogen: C10339) according to the manufacturer protocols. Slides were incubated for 3 hours in 37° C with polyclonal anti-TRF2 antibody, washed four times in 1x PBS for 5 min each and then incubated one hour at 37° C with goat polyclonal anti-rabbit Alexa Fluor 488. First and secondary antibodies were diluted in PBS supplemented with 5% FCS. Before mounting, slides were washed again as described above and dehydrated in a graded series (70, 90, 100%) of ethanol solutions for 2 min each. Slides were mounted with Prolong Gold (Life Technologies) and imaged using super resolution microscopy as described below.

### FANCD2 labelling

72 hours after transfection of the chromobody plasmids and 24 hours after addition of APH, cells were fixed in 3.7% formaldehyde/1x PBS for 10 min, then permeabilised with 0.5% Triton/1x PBS and blocked with 5% Bovine Serum Albumin (Sigma-Aldrich)/1x PBS. Slides were incubated for 3 hours at 37° C with a polyclonal rabbit anti-FANCD2 antibody, washed four x 5 min in 1x PBS, then incubated for one hour at 37° C with goat polyclonal anti-rabbit Alexa Fluor 647. First and secondary antibodies were diluted in 1x PBS supplemented with 5% FCS. Before mounting, slides were washed again as described above and dehydrated in a graded series (70, 90, 100%) of ethanol solutions for two min each. Slides were then mounted with Prolong Gold (Life Technologies) and imaged using super resolution microscopy as described below.

### Super resolution imaging

Super resolution imaging was performed on a ZEISS LSM 880 AxioObserver confocal fluorescent microscope fitted with an Airyscan detector using a Plan-Apochromat 63x 1.4 NA M27 oil objective. Cells were imaged using 1.9% excitation power of 568 nm laser, 2% excitation power of 488 laser, and 1.8% excitation power of 647 nm laser, with 1×1 binning for all laser conditions in combination with the appropriate filter sets. Ten or more z stacks (167 nm) were captured with frame scanning mode and unidirectional scanning. Z-stacks were Airyscan processed using batch mode in Zen software.

### siRNA transfection

Control non-targeting (control siRNA, D-001810-10), ATR (L-003202-10), and mTOR (L-003008-10), ON-TARGETplus siRNA pools (Dharmacon), and IPMK (D-006740-05) siGENOME individual siRNAs (Dharmacon) were transfected using Lipofectamine RNA imax (Thermofisher Scientific) according to manufacturer protocol.

### Western blotting

Preparation of whole cell extracts and western blots were done as described previously ^37^ and luminescence was visualised on an LAS 4000 Imager (Fujifilm).

### Visualising genome instability

Cells were grown on sterile glass coverslips and treated with LatB ± APH for 24 hours. Cells were then fixed in 3.7% formaldehyde/1x PBS for 10 min and permeabilised with 0.5% Triton/1x PBS and stained with 1 μg/ml DAPI (Sigma-Aldrich). Slides were mounted with Prolong Gold (Life Technologies) and imaged using Zen software and a ZEISS AxioImager Z.2, with a 63x 1.4 NA oil objective, appropriate filter cubes and an Axiocam 506 monochromatic camera. Images were scored by eye.

### Volume and ellipticity measurements

Imaging data was imported into Imaris 8.4.1 software (Bitplane AG, Zurich, Switzerland), where nuclei were segmented using the ‘Surfaces’ function on the actin-NLS channel. Volume and ellipticity were calculated via the ‘Volume’ and ‘Ellipticity’ functions. For live cell imaging, surfaces were tracked over time and volume and ellipticity were calculated for every time point.

### Nuclear localisation of replication foci and telomeres

Imaging data was imported into Imaris 8.4.1 software, where nuclei were segmented as “cells” using the Cell function on the actin-NLS channel. Telomeres or PCNA foci were segmented as “vesicles” using the Cell function. The distance between foci and the nuclear periphery were calculated using the function ‘Distance of vesicles to cell membrane’.

### Measurement of late S-phase NLS-GFP-Actin-CB foci speed and displacement

Imaging data was imported into Imaris 8.4.1 software. First, nuclei were segmented as ‘Surfaces’ using the Surface function on the actin-NLS channel. Nuclei speed was calculated using the ‘speed’ function. Then, late S-phase NLS-GFP-Actin-CB foci were segmented using the Spots function and their speed was determined using the ‘speed’ function. We subtracted nucleus speed from the foci speed to correct for cell movement. The direction of movement was determined by calculating the distance of late S-phase NLS-GFP-Actin-CB foci from the nuclear periphery in the Cell function as described above, and then subtracting the distance in the last frame from the distance in the first frame for each focus.

### Image processing for MSD analysis

Imaging data were imported into Imaris 8.4.1 software, where nuclei were coarsely segmented using the ‘Surfaces’ function on the actin-NLS channel. Cells migrate on the substrate during acquisition; this must be corrected for to accurately track the relative position of the replication foci within the nucleus over time. We therefore developed an image processing method such that the position of the foci could be quantified in the nucleus-frame-of-reference and not the laboratory-frame-of-reference (Video 6). We termed this technique “registration”. To register nuclei, positional and morphological data on the segmented nucleus was transferred into Matlab (MathWorks, Nattick, NA, USA) via the XT module of Imaris, where the centre of mass and the angle of the long axis of the 3D ellipsoid fit of the Surface were retrieved via custom Matlab code. The algorithm then placed via the XT module in Imaris Reference Frames centred on the centre of mass of the nuclei and oriented in the direction of the ellipsoid long axis at every timepoint. Only rotations in the XY plane were allowed, due to the adherent nature of the studied cells. Combined, both the translational and rotational movements of the cells could be stabilised, without the need for considering a subset of foci as immobile fiducial markers. The registration operation was completed within Imaris via the Align Image function without interpolation, allowing for rotations and retaining the same dataset size as prior to registration. Subsequently, in the registered dataset, replication foci could readily be tracked via the Spots function in Imaris and Autoregressive Motion tracking, the numerical positional data exported for further motion / MSD analysis (see below). Custom Matlab code is available upon request and will be published via public repositories when manuscript is accepted.

### Replication foci MSD analysis

Displacements for a given replication focus were extracted from its positional time-series data (its “track”). Displacement values for a given time interval were extracted from all possible temporal points in the track, i.e., are not constrained to start at time zero only. Timepoints in a replication focus’ track where positions could not be ascertained are omitted from the MSD analysis. Time intervals used in the MSD analysis comprise all multiples of the temporal gap (or temporal sampling frequency) between timeframes for the given experiment, up to a maximum value of 20% of the longest duration track in the experiment. Where the MSD analysis was performed over an entire experiment, displacements for each given time interval were pooled from all tracks before taking the mean. MSD slopes represent the gradient of a linear regression model fitted to log-log transformed mean squared displacement-time interval data.

MSD analysis was performed over individual tracks using a sliding window to provide a time-series of MSD slopes. The window width of nine positional timeframes defined the sliding window, and the MSD slope was assigned to the center (5^th^) timeframe. MSD slopes were categorised into “MSD regimes” based on: confined motion (MSD slope < 0.8), diffusive (Brownian) motion (0.8 < MSD slope < 1.2), hyperdiffusive motion (MSD slope > 1.2) and sustained hyperdiffusive motion (MSD slope > 1.2 for five or more consecutive timeframes). Experimental conditions were contrasted by pooling instances of observed MSD regimes from all tracks in each condition. Statistical significance between conditions in Figure 3C was calculated using the Python scipy.stats module’s chi2_contingency function, operating over the absolute number of timeframes each MSD regime was observed under each condition

### Antibodies

Primary antibodies used in this study: mouse monoclonal anti-BrdU (BD Biosciences: 347580, 1:5) for IdU detection, rat monoclonal anti-BrdU (abcam: ab6326, 1:25) for CldU detection, rabbit polyclonal anti-TRF2 (Novus: NB110-57130, 1:250), rabbit polyclonal anti-FANCD2 (abcam: ab2187, 1:250), rabbit monoclonal anti-Lamin A/C (Sigma-Aldrich: L1293, 1:1000), rabbit polyclonal anti-alpha tubulin (abcam: ab18251, 1:5000), rabbit polyclonal anti-H2A (abcam: ab18255, 1:2000), mouse monoclonal anti-vinculin (Sigma-Aldrich: V9131, 1:5000), rabbit polyclonal antibodies: anti-mTOR, phospho-mTOR (ser2448), phospho-P70S6 kinase (ser371) and phospho-4E-BP1 (Thr37/46) are from the mTOR substrates antibody sampler kit (Cell Signaling Technology: 9862, 1:1000), rabbit polyclonal anti-IPMK (OriGene: TA308405, 1:1000), mouse monoclonal anti-CHK1 (Cell Signaling Technology: 2360, 1:1000), rabbit monoclonal anti-phospho-CHK1 (ser345) (Cell Signaling Technology: 2348, 1:1000), rabbit monoclonal anti-RAD17 (Cell Signaling Technology: 8561, 1:1000), rabbit monoclonal anti-phospho-RAD17 (ser645) (Cell Signaling Technology:6981, 1:1000), rabbit polyclonal anti-RPA32/RPA2 (Cell Signaling Technology: 52448, 1:1000) and rabbit polyclonal anti-phospho-RPA32/RPA2 (Ser8) (Cell Signaling Technology: 83745, 1:1000). Secondary antibodies used in this study: goat polyclonal anti-mouse Alexa Fluor 488 (ThermoFisher scientific: A28175, 1:25 for DNA fibre staining and 1:1000 for fixed nuclei staining), goat polyclonal anti-rat Alexa Fluor 594 (ThermoFisher scientific: A-11007, 1:25), goat polyclonal anti-rabbit Alexa Fluor 488 (ThermoFisher scientific: A11034, 1:1000), goat polyclonal anti-rabbit Alexa Fluor 568 (ThermoFisher scientific: A-11011, 1:750), goat polyclonal anti-rabbit Alexa Fluor 647 (ThermoFisher scientific: A21245, 1:750), goat polyclonal anti-rabbit HRP (Dako: P0448, 1:1000 for anti-phospho first antibodies, otherwise 1:3000), goat polyclonal anti-mouse HRP (Dako: P0447, 1:1000 for anti-phospho first antibodies, otherwise 1:3000).

### Statistics and figure preparation

Except for the MSD analysis described above, statistical analyses were performed using GraphPad Prism. In Tukey box plots the box extends from the 25th to the 75th percentile data points and the line represents the median. The upper whisker represents data points ranging up to the 75th percentile + (1.5 × the inner quartile range), or the largest value data point if no data points are outside this range. The lower whisker represents data points ranging down to the 25th percentile – (1.5 × the inner quartile range), or the smallest data point if no data points are outside this range. Data points outside these ranges are shown as individual points. Figure legends describe the error bars, statistical methods and n for all experiments. Figures were prepared using Adobe Photoshop and Illustrator.

## ACKNOWLEDGMENTS

We thank Scott Page and the Australian Cancer Research Foundation Telomere Analysis Centre at the Children’s Medical Research Institute (CMRI) for microscopy infrastructure. David Croucher and Vihanda Wickramshinghe are thanked for their feedback. M.B. acknowledges Bitplane AG for an Imaris Developer license. N.L. is supported by a Hebrew University Smorgon Foundation Fellowship, a fellowship from the Cancer Institute NSW, and bridging funds from University of Sydney. V.P.M. is supported by an Australian Post-Graduate Award from the University of Sydney. M.N.R. is supported by the University of Sydney Centre for Excellence in Advanced Food Enginomics. M.B. is supported by funding via European Molecular Biology Laboratory (EMBL) Australia. A.J.C. is supported by grants from the Australian NHMRC (1053195, 1106241, 1104461), the Cancer Council NSW (RG 15–12), the Cancer Institute NSW (11/FRL/5–02) and philanthropy from Stanford Brown, Inc (Sydney, Australia).

## AUTHOR CONTRIBUTIONS

N.L. and A.J.C. conceived of the study. N.L. and V.P.M. performed experimentation. M.N.R. and M.B. developed novel analysis tools. N.L., M.N.R., M.B., and A.J.C. analysed the data. N.L., M.B., and A.J.C created figures and wrote the manuscript.

## AUTHOR INFORMATION

All data are archived at CMRI or the University of New South Wales. The authors declare no competing interests. The chromobodies used in this study were purchased with an MTA and can be attained from Chromotek. All data is available in the main text, extended data and videos. Custom code for live cell imaging analysis is currently available upon request from MB and will be made available via public repositories (GitHub and Imaris Open) prior to publication. Correspondence and requests for materials should be addressed to A.J.C.

---

**Extended Data Fig. 1:**
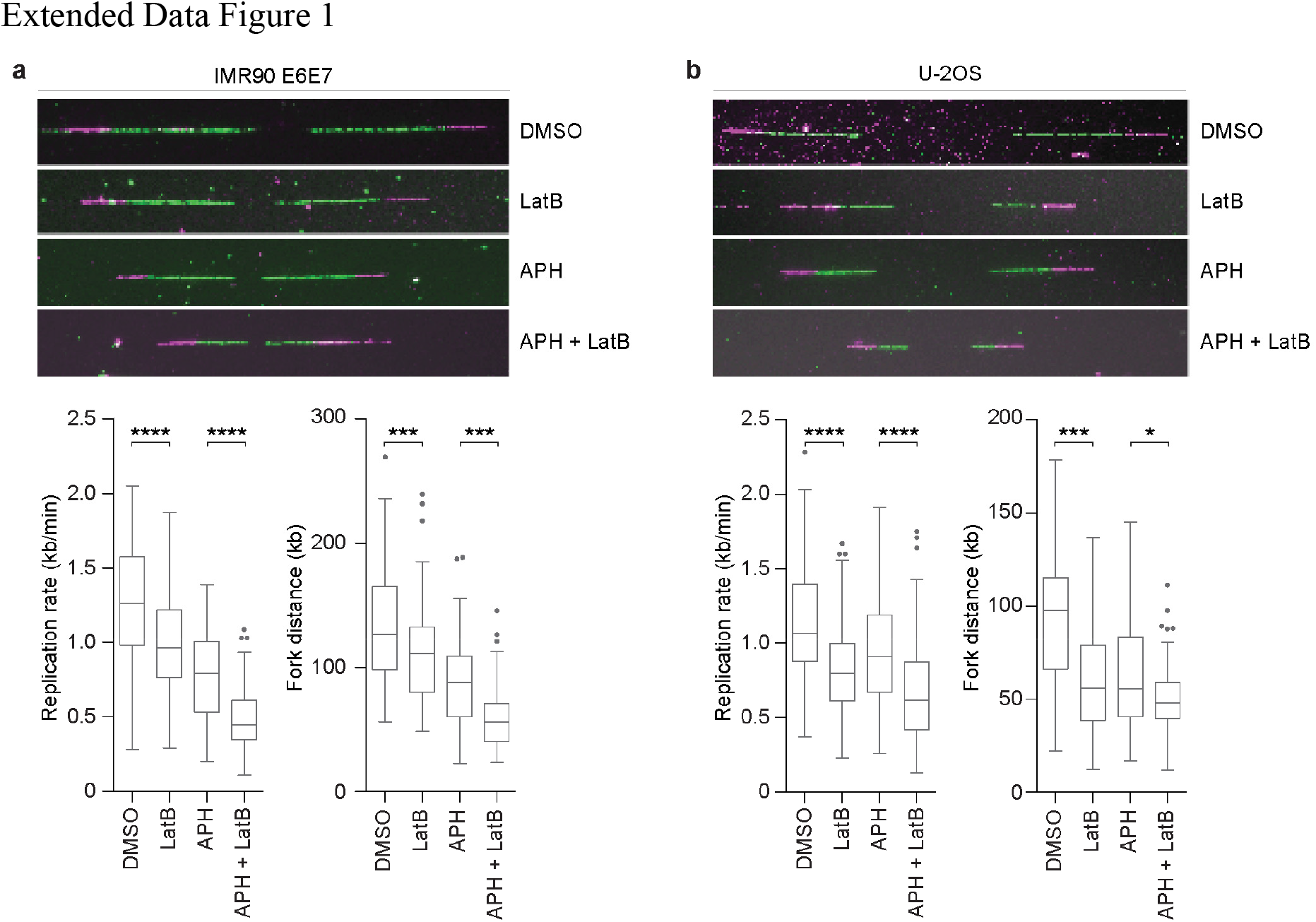
Inhibiting actin dynamics decreases replication rate and fork distance. **a-b,** IMR90 E6E7 (**a**) and U-2OS (**b**) cultures were treated with 0.01 μM Aphidicolin (APH) and/or 0.2 μM Latrunculin B (LatB) for two hours (one hour before and for one hour during pulse-labelling with thymidine analogous) and replication dynamics measured by chromatin fibre assays. Representative images are shown above, and the calculated replication rate and fork distance are shown below (n ≥ 110 forks per condition for IMR90 E6E7 and n ≥ 116 forks per condition for U-2OS are compiled into a Tukey box plot, two-tailed student’s t-test, *\*p < 0.05, ***p < 0.0005, ****p < 0.0001*.

**Extended Data Fig. 2:**
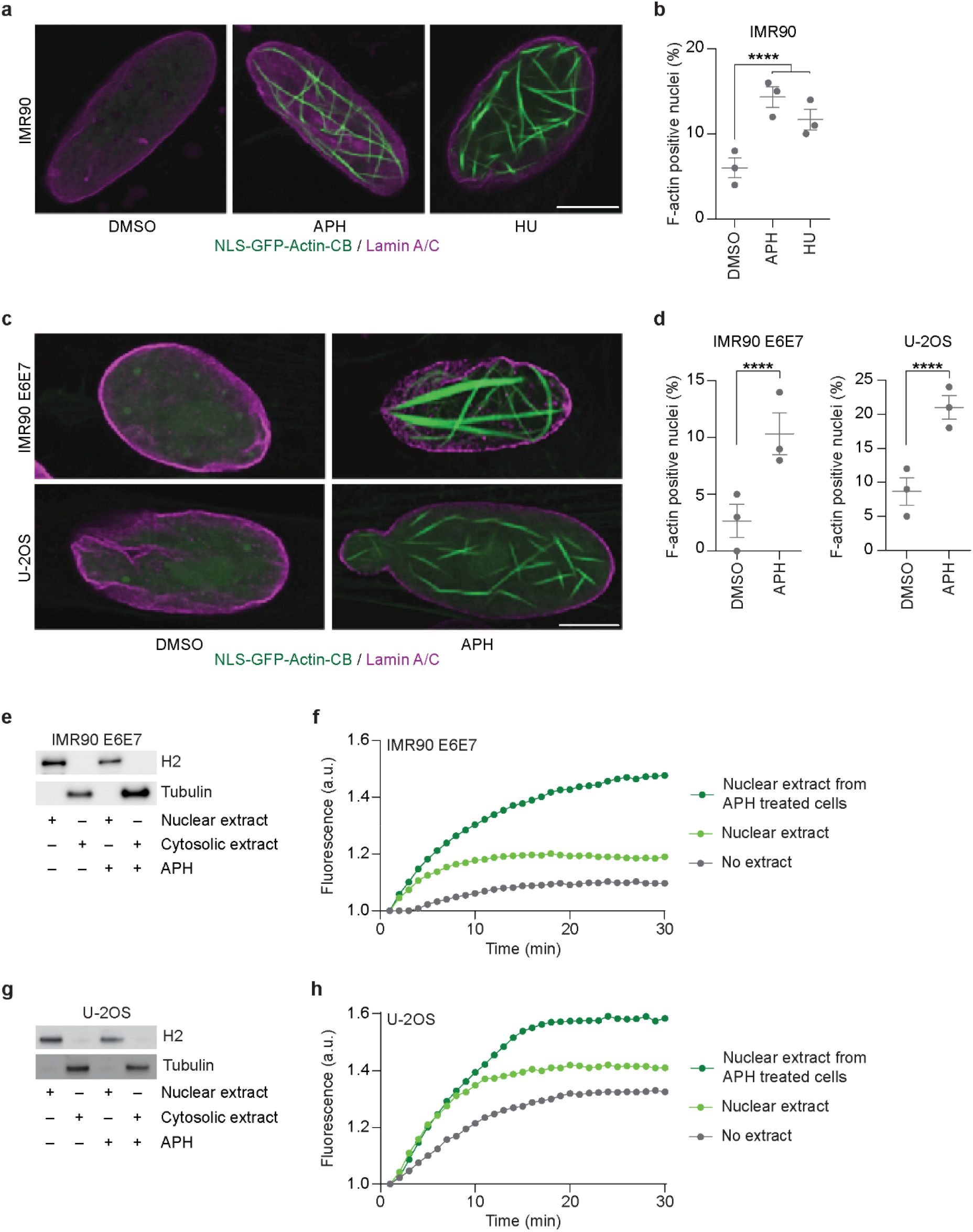
Replication stress induces nuclear actin polymerisation. **a-d,** NLS-GFP-actin-CB transfected IMR90 (**a, b**) IMR90 E6E7 (**c, d**), and U-2OS (**c, d**) cultures were treated with 0.4 μM APH or 500 μM HU for eight hours before fixation and staining for Lamin A/C. Representative images are from a single Z-plane through the nuclear volume captured by super-resolution microscopy. Quantitative data are mean ± s.e.m, with each data point representing the mean of an individual biological replicate (n ≥ 294 nuclei for IMR90, n ≥ 285 nuclei for IMR90 E6E7, and n ≥ 273 nuclei for U-2OS over three biological replicates, Fisher’s exact test). **e,** Immunoblot of nuclear and cytosolic extracts from IMR90 E6E7 fibroblasts ± 0.4 μM APH for eight hours. **f,** Representative normalised time course of pyrene-labelled actin assembly in the presence or absence of IMR90 E6E7 nuclear extracts from e. One of three independent biological replicates are shown. **g,** Immunoblot of nuclear and cytosolic extracts from U-2OS cells ± 0.4 μM APH for eight hours. **h,** Representative normalised time course of pyrene-labelled actin assembly in the presence or absence of U-2OS nuclear extracts from **g.** One of three independent biological replicates are shown. For all panels: scale bar represents 5 μm, *\*\*\*\*p < 0.0001*.

**Extended Data Fig. 3:**
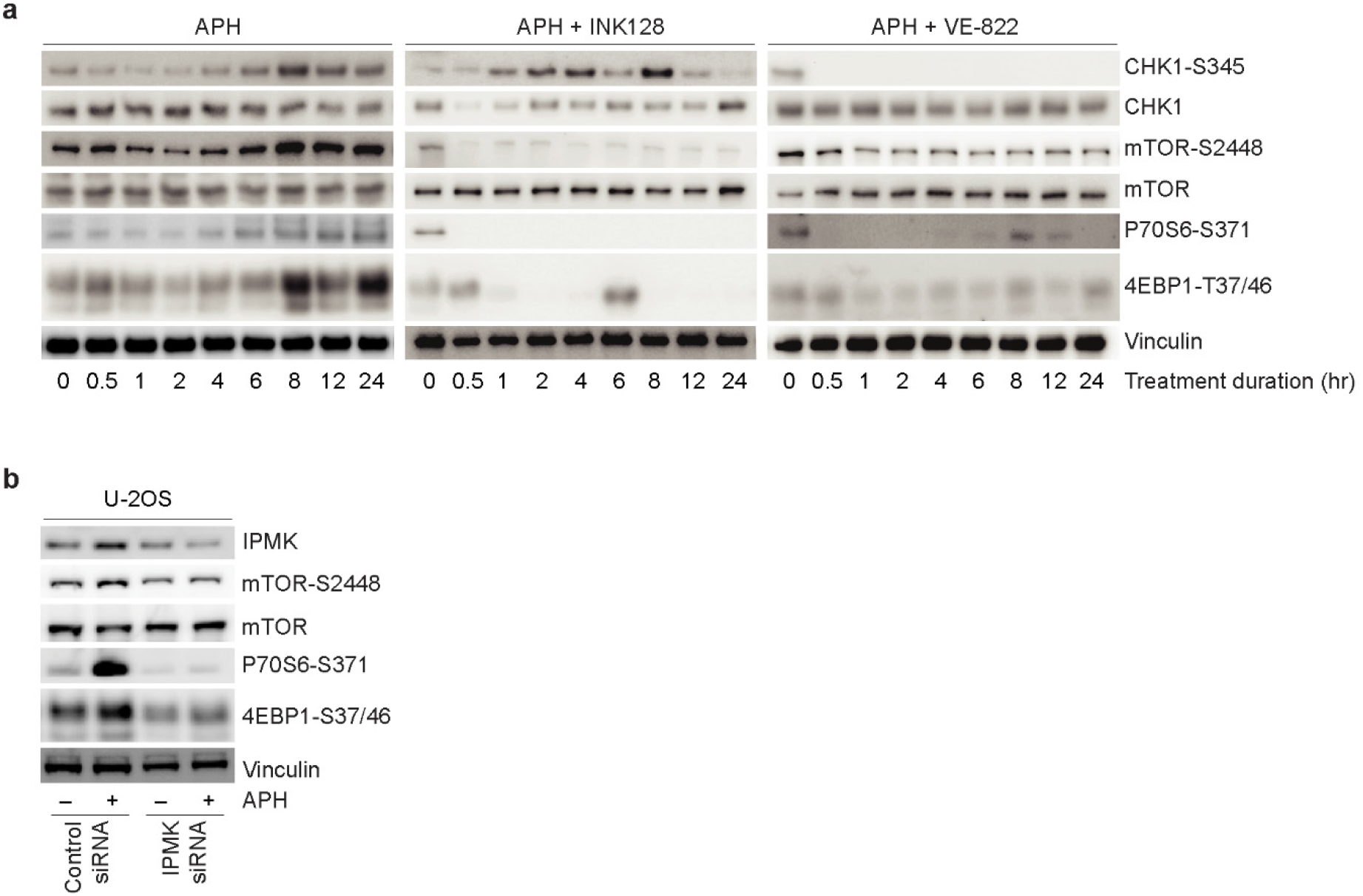
Replication stress activates mTOR signaling in an ATR- and IPMK-dependent manner. **a,** Western blots of whole cell extracts from IMR90 E6E7 cells treated with 0.4 μM APH ± 0.2 μM INK128 or 1 μM VE822 for the indicated duration. **b,** Western blots of whole cell extracts from siRNA transfected U-2OS cells treated with 0.4 μM APH for eight hours. Cells were siRNA transfected 48 hours prior to extraction.

**Extended Data Fig. 4:**
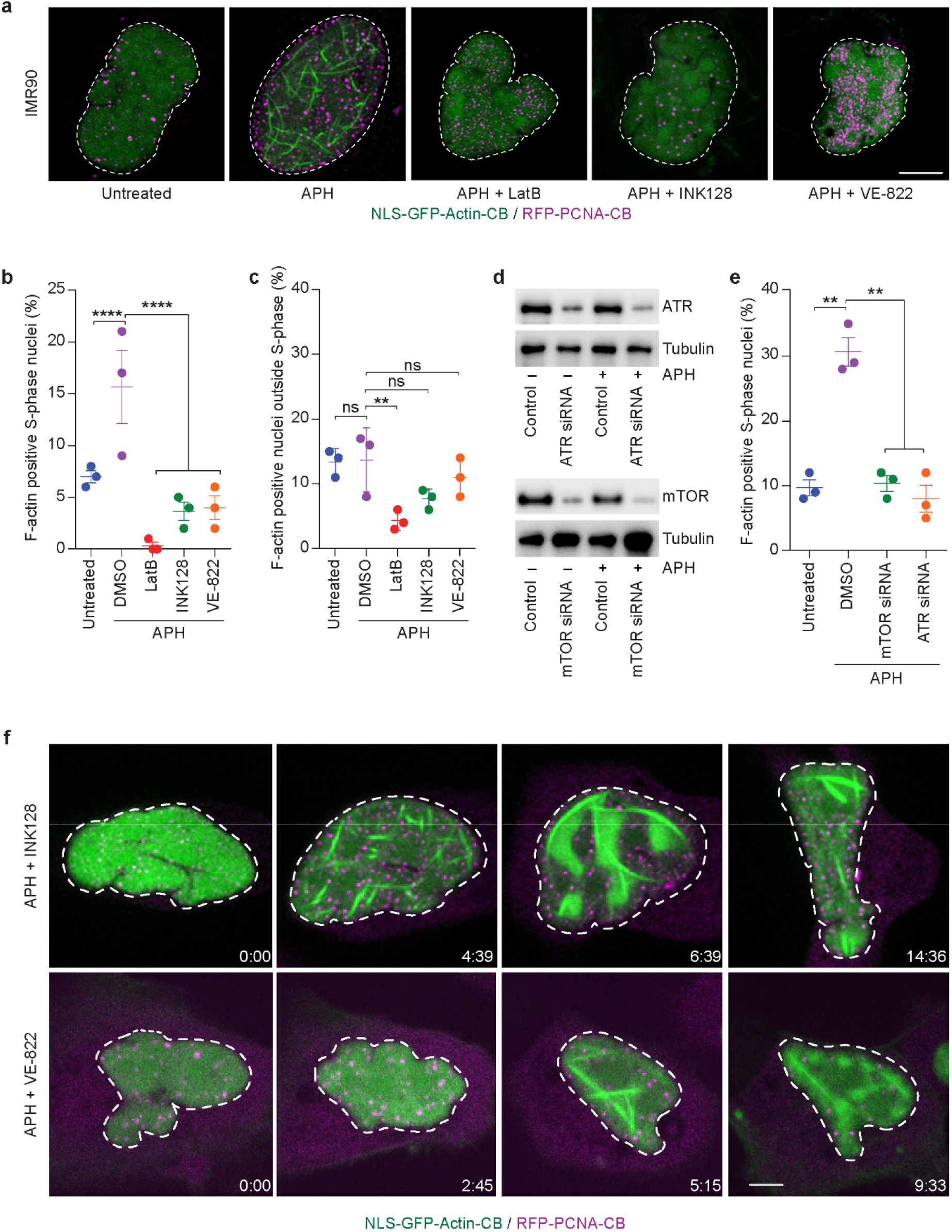
ATR and mTOR regulate F-actin to change nuclear architecture in response to replication stress. **a,** Single Z-planes from super-resolution microscopy of fixed NLS-GFP-Actin-CB and RFP-PCNA-CB transfected IMR90 fibroblasts treated with 0.4 μM APH ± 0.2 μM LatB, 0.2 μM INK128 or 1 μM VE822 for 24 hours. **b-c,** Frequency of nuclear actin fibre positive S-phase (**b**) and non-S-phase (**c**) nuclei from the experiment shown in a (mean ± s.e.m., each data point represents the mean of an individual biological replicate, n ≥ 185 and > 142 respectively for S-phase and non-S-phase nuclei over three biological replicates, Fisher’s exact test). **d,** Western blots of whole cell extracts from siRNA transfected U-2OS cells ± 0.4 μM APH for 8 hours. Cells were siRNA transfected 72 hours prior to extraction. **e,** Frequency of nuclear actin fibre positive S-phase nuclei in U-2OS cells from the time points in **d** (mean ± s.e.m., each data point represents the mean of an individual biological replicate, n ≥ 78 S-phase nuclei over three biological replicates, Fisher’s exact test). **f,** Representative images from spinning-disk live cell microscopy of NLS-GFP-Actin-CB and RFP-PCNA-CB transfected U-2OS cells treated with 0.4 μM APH + 0.2 μM INK128 or 1 μM VE822. Time is shown as (hr:min) relative to the first image of the series. For all panels: scale bar represents 5 μm, ns = not significant, *\*\*p < 0.01, ****p < 0.0001*.

**Extended Data Fig. 5:**
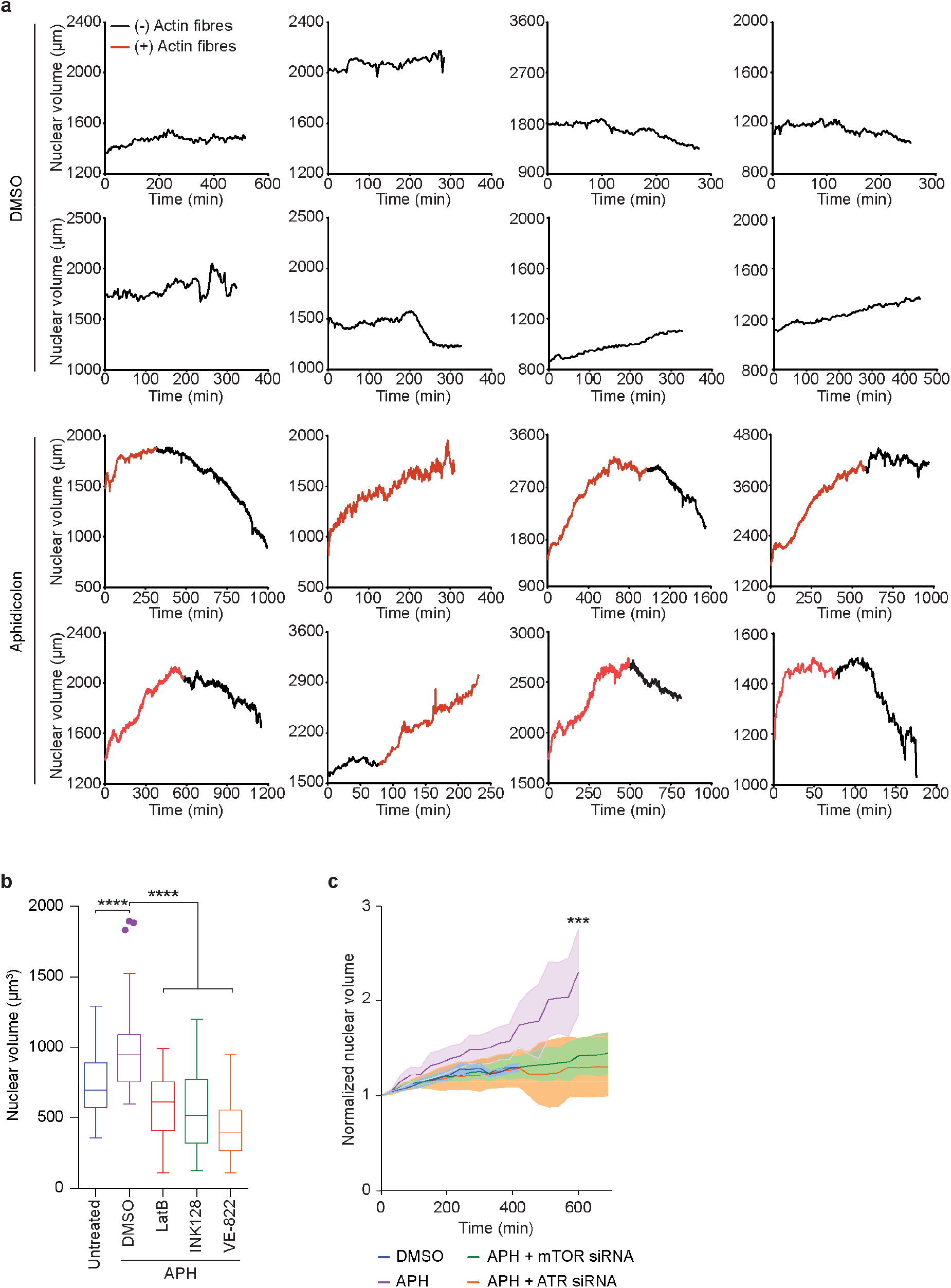
F-actin dynamics change nuclear volume in response to replication stress. **a,** Time courses from spinning disk confocal live cell imaging depicting nuclear volume of individual S-phase U-2OS cells transfected with the NLS-GFP-actin-CB and RFP-PCNA-CB in the presence or absence of 0.4 μM APH. Red line colouration represents when nuclear F-actin fibres were visible. **b,** S-phase nuclear volume from fixed NLS-GFP-actin-CB and RFP-PCNA-CB transfected IMR90 cells (n >124 cells over three biological replicates compiled into a Tukey box plot, two-tailed student’s t-test). **c,** Time courses of normalised S-phase nuclear volume from spinning disk confocal live cell imaging of NLS-GFP-actin-CB and RFP-PCNA-CB transfected U-2OS cells, 42-96 hours post transfection with the indicated siRNAs, in cultures ± 0.4 μM APH (mean ± s.e.m., n ≥ 24 nuclei over five biological replicates, one-way ANOVA). For all panels: *\*\*\*p < 0.001, ****p < 0.0001*.

**Extended Data Fig. 6:**
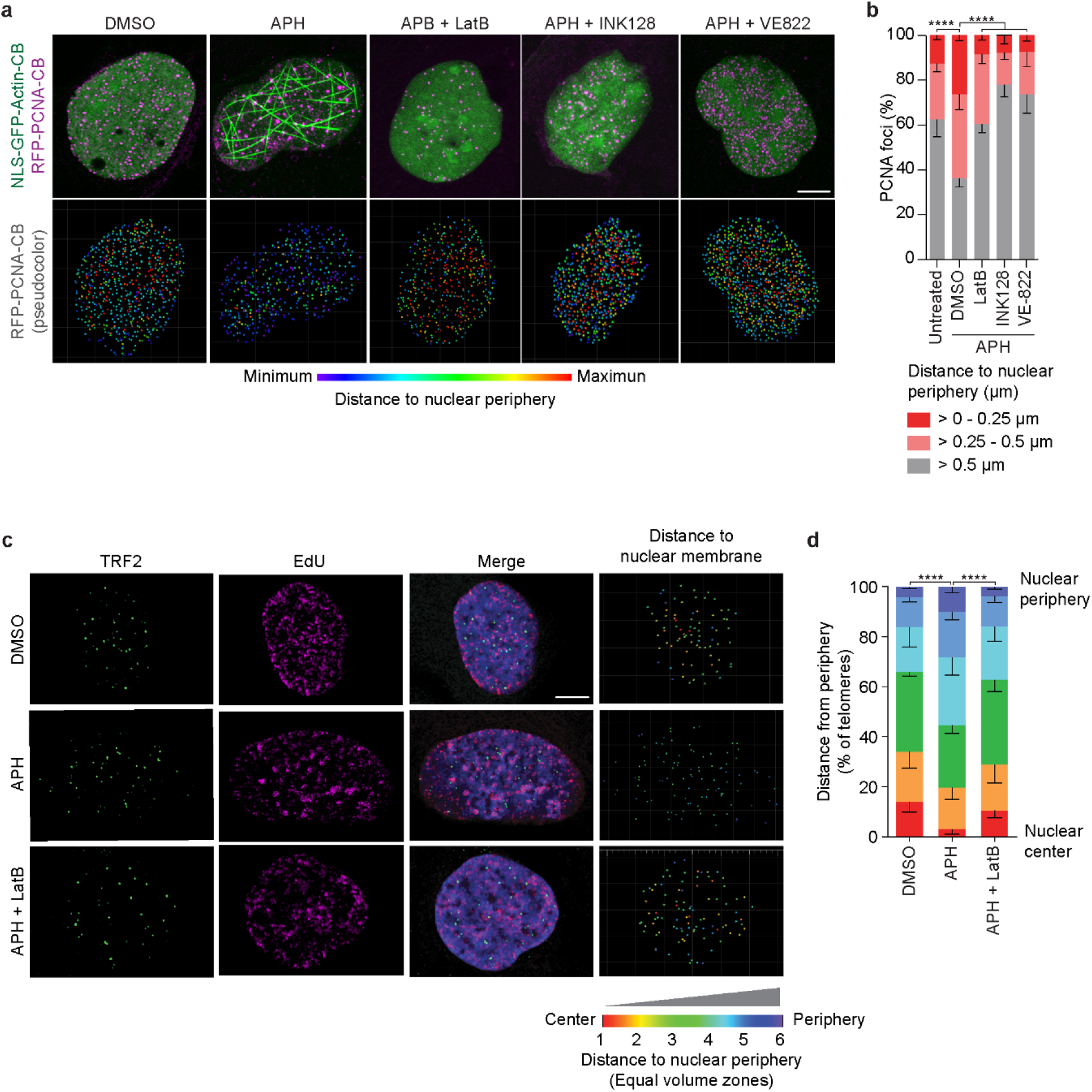
Actin polymerisation mobilises replication foci in cells experiencing replication stress. **a,** Upper panel: Representative super-resolution microscopy of a single Z-plane taken from a three-dimensional image through the nuclear volume of fixed NLS-GFP-actin-CB and RFP-PCNA-CB transfected IMR90 cells treated with 0.4 μM APH ± 0.2 μM LatB, 0.2 μM INK128 or 1 μM VE822 for 24 hours. Lower panel: RFP-PCNA-CB foci were identified throughout the three-dimensional volume of the above images and their distances from nuclear periphery identified. All foci from the three-dimensional image shown above are collapsed into two dimensions and the distance from the nuclear periphery identified via colour coding. **b,** Distance of RFP-PCNA-CB foci from nuclear periphery from the experiments depicted in **a.** (mean ± s.e.m., n ≥1684 foci from at least 12 nuclei, chi square test). **c,** Representative super-resolution microscopy of a single Z-plane taken from a three-dimensional image throughout the nuclear volume of fixed U-2OS cells treated with 0.4 μM APH ± 0.2 μM LatB. Cells were labelled with EdU for one hour prior to fixation to identify S-phase cells and stained with a TRF2 antibody. Telomeres were identified throughout the nuclear volume of S-phase cells and their distance to the nuclear periphery calculated. Nuclei were segmented into six equal volume zones from nuclear center to the nuclear periphery and each telomere assigned to the corresponding zone. All telomere foci from the three-dimensional image are collapsed into two dimensions in the far-right panel, and the zone for each telomere relative to the nuclear periphery is identified via colour coding. **d,** Distance of individual telomeres to nuclear periphery from the images depicted in **c.** (mean ± s.e.m., n ≥ 2096 telomeres from at least 29 nuclei, chi square test). For all panels: Scale bar represents 5 μm, *\*\*\*\*p < 0.0001*.

**Extended Data Fig. 7:**
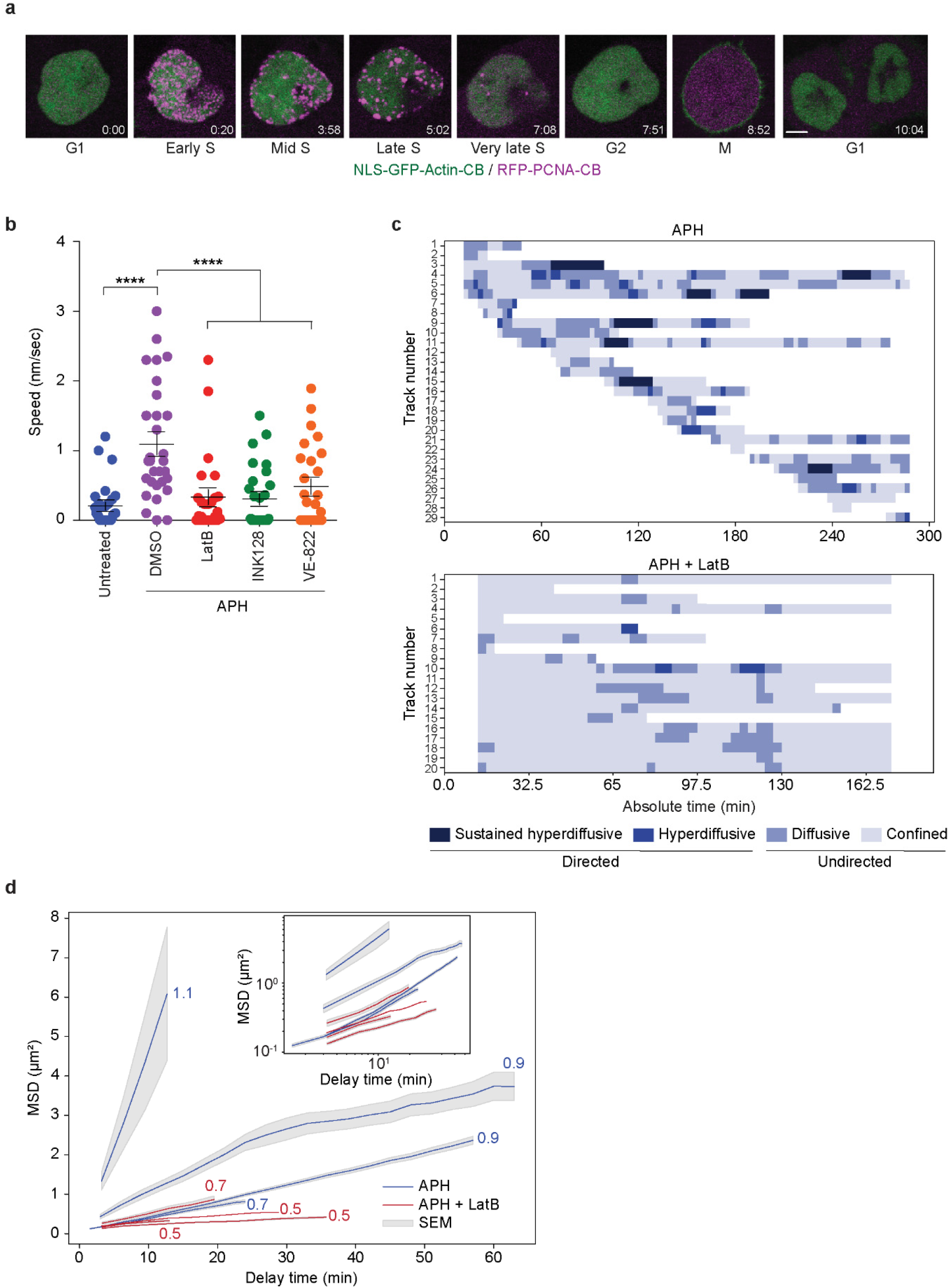
F-actin facilitates directed movement of late replication foci towards nuclear periphery. **a.** Representative images of a complete cell cycle taken by spinning-disk live cell microscopy of a NLS-GFP-Actin-CB and RFP-PCNA-CB transfected U-2OS cell. Time is shown as (hr:min) relative to the first image of the series. Scale bar represents 5 μm. **b.** Speed of late S-phase RFP-PCNA-CB foci in U-2OS cells treated with 0.4 μM APH ± 0.2 μM LatB, 0.2 μM INK128 or 1 μM VE822 (mean ± s.e.m., n ≥ 26 late foci from at least 7 nuclei, two-tailed student’s t-test, *\*\*p < 0.005, **** p < 0.0001*). **c,** Representative late S-phase RFP-PCNA-CB foci trajectories from U-2OS treated with 0.4 μM APH ± 0.2 μM LatB. Data are displayed as heat-maps indicating the regime of movement based on MSD analysis throughout the lifetime (in absolute min) of the individual tracks. **d,** Mean Squared Displacement (MSD) analysis of replication foci trajectories in NLS-GFP-Actin-CB and RFP-PCNA-CB transfected U-2OS cells treated with 0.4 μM APH ± 0.2 μM LatB, 0.2 μM INK128 or 1 μM VE822. Inset shows MSD curves in log-log space, used to calculate linear regression slopes independently for each replicate experiment. Slope values are reported alongside each series in the main panel (n ≥ 45 tracks from four nuclei for each condition).

**Extended Data Fig. 8:**
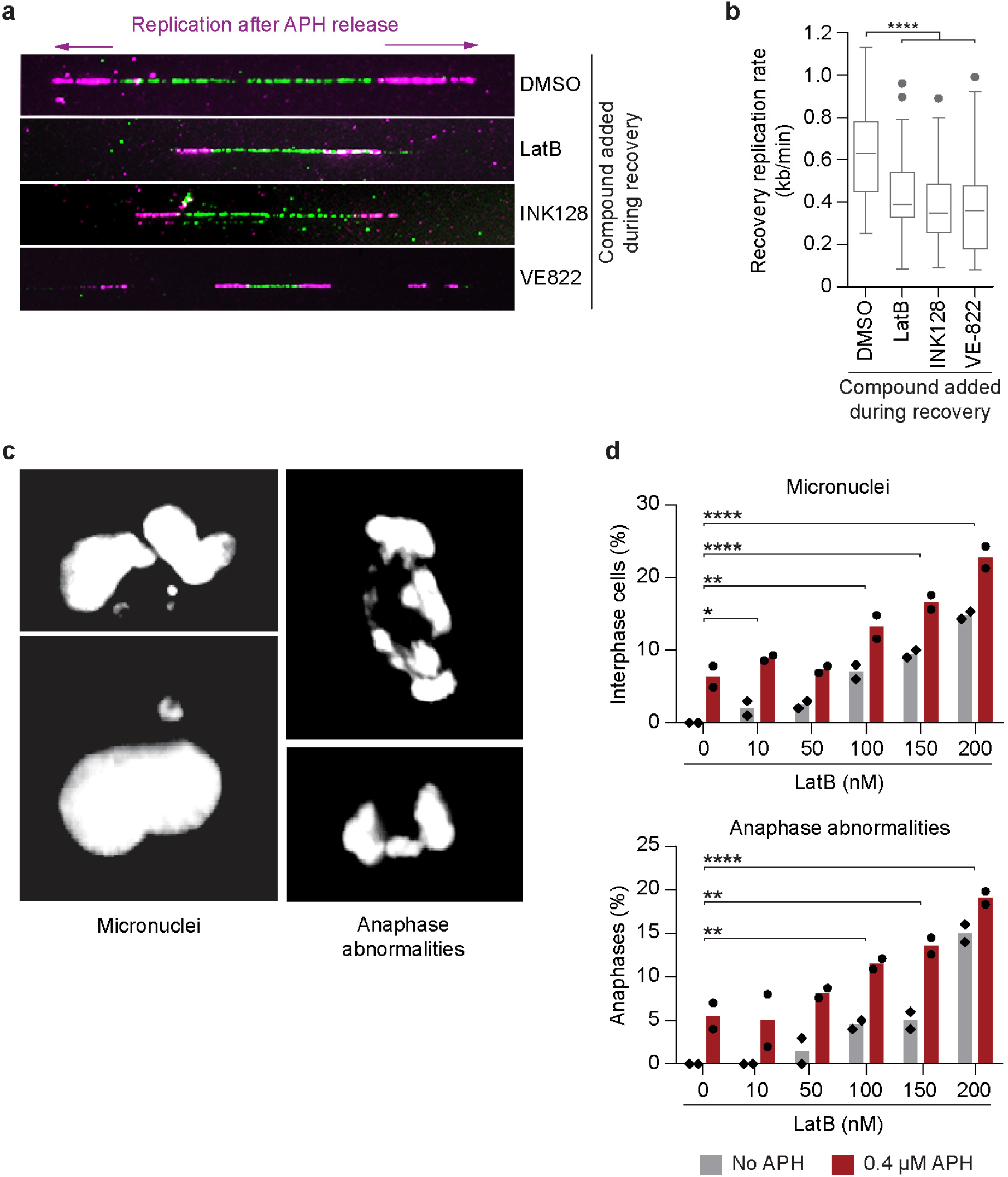
Actin dynamics mediate replication fork repair to maintain genome stability. **a,** Representative images of chromatin fibre assays to measure replication fork restart in IMR90 E6E7 cells. Experiments were performed as depicted in Figure 4a. Cells were treated with 0.1 μM APH and IdU for three hours, before washout and labelling with CldU ± 0.2 μM LatB, 0.2 μM INK128, or 1 μM VE-822. Replication rate is measured from the CldU track. **b,** Quantitation of the experiment shown in **a** (n > 132 replication forks from two biological replicates are compiled into a Tukey box plot, two-tailed student’s t-test). **c,** Representative images of micronuclei (left panels) and anaphase abnormalities (right panels) in IMR90 E6E7 cells following treatment with escalating dosages of LatB. **d,** Frequency of micronuclei (upper panel) and anaphase abnormalities (bottom panel) from the images depicted in c (n ≥ 158 cells for each concentration over two biological replicates, individual replicate means are shown by the data points, bar represents the overall mean. Fisher’s exact test). For all panels, *\*p < 0.01, **p < 0.001, ***p < 0.0005, ****p < 0.0001*.

**Video 1: Normal cell cycle in U-2OS cells expressing the NLS-GFP-actin and RFP-PCNA chromobodies.** Representative maximum projection movie captured by spinning-disk confocal live cell microscopy of U-2OS cells transfected with the NLS-GFP-actin-CB and RFP-PCNA-CB. Cells were transfected 72 hours before imaging and treated with DMSO starting 24 hours before the imaging session. Time is shown as (hr:min:sec) relative to the first image of the series.

**Video 2: Replication stress induces nuclear F-actin.** Representative maximum projection movie captured by spinning-disk confocal live cell microscopy of a U-2OS cells transfected with the NLS-GFP-actin-CB and RFP-PCNA-CB and treated with 0.4 μM APH. Cultures were transfected 72 hours before imaging and treated with APH starting 24 hours before the imaging session. Time is shown as (hr:min:sec) relative to the first image of the series.

**Video 3: LatB prevents nuclear actin polymerisation in APH treated cells.** Representative maximum projection movie captured by spinning-disk confocal live cell microscopy of a U-2OS cell transfected with the NLS-GFP-actin-CB and RFP-PCNA-CB and treated with 0.4 μM APH and 0.2 μM LatB. Cultures were transfected 72 hours before imaging and treated with APH and LatB starting 24 hours before the imaging session. Time is shown as (hr:min:sec) relative to the first image of the series.

**Video 4: Inhibiting mTOR or ATR prevents nuclear actin polymerisation in APH treated cells.** Representative maximum projection movie captured by spinning-disk confocal live cell microscopy of a U-2OS cell transfected with the NLS-GFP-actin-CB and RFP-PCNA-CB and treated with 0.4 μM APH and 0.2 μM INK128 or with 0.4 μM APH and 1 μM VE-822. Cultures were transfected 72 hours before imaging and treated with APH and INK128 starting 24 hours before the imaging session. Time is shown as (hr:min:sec) relative to the first image of the series.

**Video 5: Inhibiting mTOR results in loss of nuclear F-actin structural integrity in cells under replication stress.** Experimental conditions are identical to the cell treated with 0.4 μM APH and 0.2 μM INK128 in Video 4. The F-actin structural collapse phenotype is demonstrated.

**Video 6: Demonstration of nuclear registration for RFP-PCNA-CB foci analysis.** The same cell is shown before and after registration. The video is a maximum projection movie captured by spinning-disk confocal live cell microscopy of a U-2OS cell transfected with the NLS-GFP-actin-CB and RFP-PCNA-CB. Cultures were transfected 72 hours before imaging. Time is shown as (hr:min:sec) relative to the first image of the series.

**Video 7: Replication stress induces diffusive movement of late replication foci.** A presentative registered maximum projection movie from spinning-disk confocal live cell microscopy of late S-phase of NLS-GFP-actin-CB and RFP-PCNA-CB transfected U-2OS cells treated with 0.4 μM APH. Experimental conditions are the same as Video 2. The nucleus is registered as shown in Video 6 and foci were tracked. Time is shown as (hr:min:sec) relative to the first image of the series.

**Video 8: Diffusive movement of stressed late replication foci is dependent on actin polymerisation.** A representative registered maximum projection movie from spinning-disk confocal live cell microscopy of late S-phase of NLS-GFP-actin-CB and RFP-PCNA-CB transfected U-2OS cells treated with 0.4 μM APH and 0.2 μM LatB. Experimental conditions are the same as Video 3. The nucleus is registered as shown in Video 6 and foci were tracked. Time is shown as (hr:min:sec) relative to the first image of the series.

**Video 9: Replication stress induces directed movement of late replication foci along actin fibres.** Experimental and analysis conditions are the same as Video 7. Directed movement of a late replication foci along a nuclear F-actin filament is shown.

